# Multi-Scale 3D Cryo-Correlative Microscopy for Vitrified Cells

**DOI:** 10.1101/2020.05.21.107771

**Authors:** Gong-Her Wu, Patrick G. Mitchell, Jesus G. Galaz-Montoya, Corey W. Hecksel, Emily M. Sontag, Vimal Gangadharan, Jeffrey Marshman, David Mankus, Margaret E. Bisher, Abigail K. R. Lytton-Jean, Judith Frydman, Kirk Czymmek, Wah Chiu

**Affiliations:** Department of Bioengineering, James H. Clark Center, Stanford University, Stanford, CA 94305; Division of CryoEM and Bioimaging, SSRL, SLAC National Accelerator Laboratory, Menlo Park, CA 94025; Department of Biology, James H. Clark Center, Stanford University, Stanford, CA 94305; Zeiss Research Microscopy Solutions, White Plains, NY 10601; David H. Koch Institute for Integrative Cancer Research, Massachusetts Institute of Technology, Cambridge, Massachusetts, 02139; Advanced Bioimaging Laboratory, Donald Danforth Plant Science Center, Saint Louis, MO 63132

**Keywords:** Airyscan microscopy, cryo correlative-light and electron microscopy, cryoCLEM, volume cryo-focused ion bean scanning electron microscopy, cryoFIB-SEM, cryo-electron tomography, cryoET, Hsp104 chaperone, protein aggregation

## Abstract

Three-dimensional (3D) visualization of vitrified cells can uncover structures of subcellular complexes without chemical fixation or staining. Here, we present a pipeline integrating three imaging modalities to visualize the same specimen at cryogenic temperature at different scales: cryo-fluorescence confocal microscopy, volume cryo-focused ion beam scanning electron microscopy, and transmission cryo-electron tomography. Our proof-of-concept benchmark revealed the 3D distribution of organelles and subcellular structures in whole heat-shocked yeast cells, including the ultrastructure of protein inclusions that recruit fluorescently-labelled chaperone Hsp104. Since our workflow efficiently integrates imaging at three different scales and can be applied to other types of cells, it could be used for large-scale phenotypic studies of frozen-hydrated specimens in a variety of healthy and diseased conditions with and without treatments.

## INTRODUCTION

Visualizing cells in 3D is a powerful approach to study structures and interactions of organelles and macromolecular complexes. Among 3D imaging techniques for cell biology, cryogenic electron tomography (cryoET) is emerging as a leading approach for *in situ* structure determination^1^. However, several limitations preclude its wider application, including the thickness of some samples and difficulties in locating and identifying features of interest within them. To visualize molecular details in thicker samples by cryoET, such as regions in eukaryotic cells away from the thin cell periphery, cryogenic focused ion beam scanning electron microscopy (cryoFIB-SEM) has been used to generate thin lamellae from vitrified cells^2-5^ (i.e. thin layers through the bulky cell) with a process called “ion beam milling”, enabling many exciting biological observations inside the cell^6^. However, technical challenges remain in ensuring that the milled lamellae contain the features of interest. Correlative light and electron microscopy (CLEM) can overcome this challenge by fluorescently labelling targets^7,8^, thereby guiding cryoFIB-SEM milling^9,10^ and the selection of optimal imaging areas for cryoET experiments, as well as aiding the interpretation of observed features. However, even the smallest eukaryotic cells are typically several microns thick and the desirable lamella thickness range is ∼100-500 nm; thus, targeting nanoscale features along the z direction remains extremely challenging. Here, as a proof-of-principle, we present a pipeline to address this challenge by using high-resolution CryoAiryscan Confocal Microscopy (CACM) to determine the z position of fluorescent targets within cells that were vitrified on electron microscopy (EM) grids, followed by cryoFIB-SEM “mill and view” (MAV) imaging, which provides a 3D view of whole cells with resolvable organelles as they are being milled to produce a lamella containing the target of interest, and ending with visualization of molecular details of regions of interest by cryoET.

## RESULTS

Our 3D multi-scale correlative imaging pipeline integrates three different microscopy platforms with different resolution ranges and contrast mechanisms, operated at cryogenic temperature (**Figure 1A,B, Supplementary Figure 1, & Supplementary Movie 1**). To test our workflow, we applied all steps to the same vitrified *Saccharomyces cerevisiae* cells (yeast, ∼5-10 μm in diameter) in which the chromosomal gene *hsp104* coding for chaperone heat shock protein Hsp104 bears an in-frame C-terminal green fluorescent protein (GFP) tag (hereafter Hsp104-GFP). Integration of such tag does not affect Hsp104 functions nor cell viability^11-13^. Upon heat stress, Hsp104-GFP is recruited to inclusions that appear as bright puncta in fluorescence microscopy images^14^.

**Figure 1.**
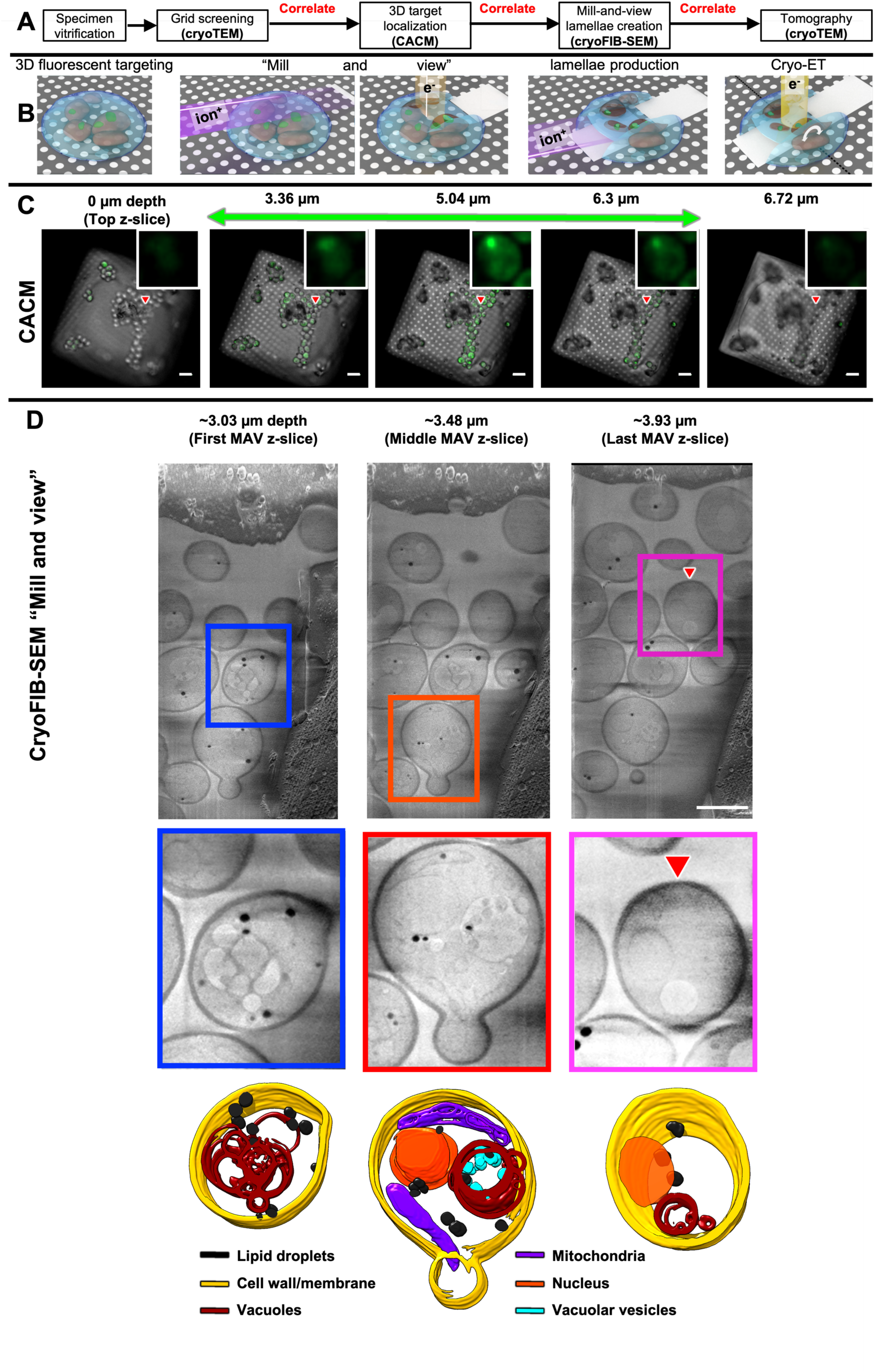
**(A)** Experimental workflow including multiple correlative imaging steps under cryogenic conditions via different imaging platforms. **(B)** Cartoon representation from left to right: vitrified yeast cells on an EM grid containing a tagged target; MAV-mode imaging by cryoFIB-SEM; lamella generation; cryoET imaging. **(C)** Representative CACM images showing slices through heat-shocked yeast cells expressing Hsp104-GFP on an EM grid, recorded at different z heights (scale bar=10 μm). The green signals come from our tagged target and the red triangle points to the location of a bright fluorescent punctum, indicative of a large inclusion containing Hsp104-GFP, highlighted in the zoomed-in insets. **(D)** Representative MAV images at different z heights (starting at ∼3.36 μm depth from the cell surface, as guided by the CACM images in (**C**)). The colored boxes highlight individual yeast cells showcasing different contextual subcellular features, annotated in the bottom row, as labelled (scale bar=5 μm), and the red triangle points to densities at the location of the same punctum singled out in (**C**).

We first imaged our vitrified specimens using CACM. To determine the relative z-height of each region of interest (ROI) containing fluorescent puncta inside the yeast cells, we counted the number of z-slices from the top of the stack to the center of each fluorescent punctum in the CACM z-stacks. The punctum shown in **Figure 1C** became visible at ∼2.94 μm depth from the top surface of the cell and persisted through an additional ∼3.36 μm. We used the correlative software ZEN Connect (Zeiss) to overlay a CACM image of the ROI with a corresponding cryoFIB-SEM image and readily located the target punctum containing Hsp104-GFP in 3D within the achievable z-resolution of the optical microscope (**Supplementary movie 2**). We used this CACM data as a guide to mill away a large portion from the top of the cell (∼3 μm) to approach the ROI indicated by the fluorescent inclusion. Then, we used the “serial milling block-face imaging”^9,15^ mode of the cryoFIB-SEM to mill 31 finer slices (∼30 nm thickness for each slice or a total of ∼930 nm) and collected an SEM image of the freshly-exposed surface in each slice, a process hereafter referred to as “mill and view” (MAV) for frozen specimens in a near-native state without exogenous heavy-metal stains. This imaging mode provided rich contextual structural information of our vitrified specimen above the region ultimately prepared as a lamella, thereby generating a 3D atlas of multiple whole yeast cells at intermediate resolution as we approached the target (**Figure 1D**). Our MAV images revealed abundant lipid droplets, vacuoles with vesicle-like features, mitochondria, cell nuclei, and daughter cells in the process of budding. Importantly, we minimized the electron beam dose during milling to optimally preserve the specimen for subsequent cryoET experiments.

The correlation between CACM (**Figure 2A**) and cryoFIB-SEM MAV (**Figure 2B**) data in 3D using Zen Connect software (Zeiss) (**Figure 2C**) allowed to identify areas on the lamella (**Figure 2D,E**) with fluorescent puncta from which to collect cryoET tilt series. We also collected tilt series from areas with diffuse or no fluorescence as internal controls. We generated ∼290-500 nm-thick lamellae to include as much of the 3D context proximal to the inclusions as permitted by the penetration power of the 300 keV transmission electron microscope. The tomograms from regions containing the bright puncta (*i.e.* Hsp104-GFP-positive inclusions) exhibited large and dense pleomorphic structures (**Figure 2F**), likely consisting of misfolded proteins. Interestingly, unlike some prior observations by traditional electron microscopy^16^, these inclusions did not appear as a continuous or completely amorphous mass but rather as a collection of pleomorphic compact granules with distinct boundaries. These granules were similar in size, and appeared to cluster in one specific region of the cell^17^, giving rise to inclusions that, with lower resolution microscopy approaches, may appear to be continuous structures.

**Figure 2.**
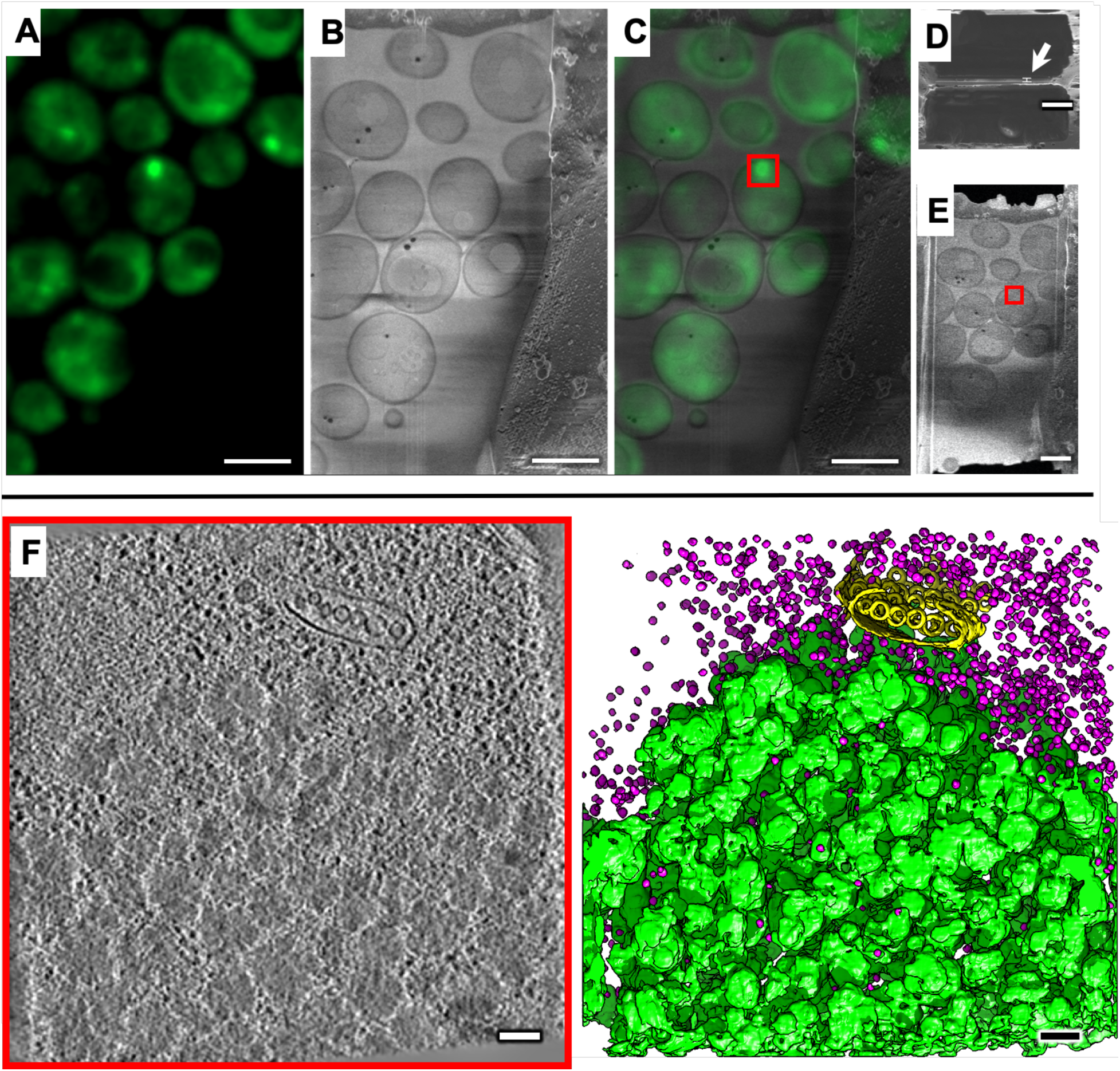
(**A).** Z-directional, maximum intensity projection of CACM z-stack images of yeast cells expressing Hsp104-GFP. **(B)** The last cryoFIB-SEM MAV slice, right before creating the lamella. **(C)** Superposition between (A) and (B) to correlate fluorescent signals with areas seen by cryoFIB-SEM to select optimal regions on the lamella that both contain the target and are amenable for subsequent cryoET experiments. **(D)** Top and **(E)** slanted views of the final milled lamella (white arrow highlights an estimated thickness of ∼260-300 nm). **(F)** Slice (∼3 nm thick) through a representative cryoET tomogram collected from a region with a bright fluorescent punctum and corresponding annotation of features (bottom row) in different colors (green:inclusions containing Hsp104-GFP; yellow:a multivesicular body made of lipid membranes; pink:ribosomes. Scale bars: A-C, 5 μm; D, 2.5 μm; E, 100 nm).

The fluorescence intensities seen by CACM (**Figure 2A**) correlated with the abundance and size of pleomorphic Hsp104-GFP-containing clusters in cryoET tomograms (**Figure 2F, Figure 3, & Supplementary movie 2**). Notably, tomograms from regions with diffuse GFP fluorescence also contained clusters made of pockets with a similar size and shape (**Figure 3A,B**) as those observed in the large inclusion (**Figure 2F**), but they were more sparsely distributed through the cytoplasm. On the other hand, tomograms from regions with no fluorescence (**Figure 3C**) or from control cells without heat shock lacked these clusters altogether. In addition to providing insight into the morphology of Hsp104-GFP clusters, the tomograms revealed ribosomes, multi-vesicular bodies (MVBs), mitochondria, vesicles budding into vacuoles, nuclei with nuclear pores, among other subcellular features, some of which we annotated semi-automatically using neural networks in EMAN2^18^ (bottom row in **Figure 2 & 3, Supplementary movie 2**).

**Figure 3.**
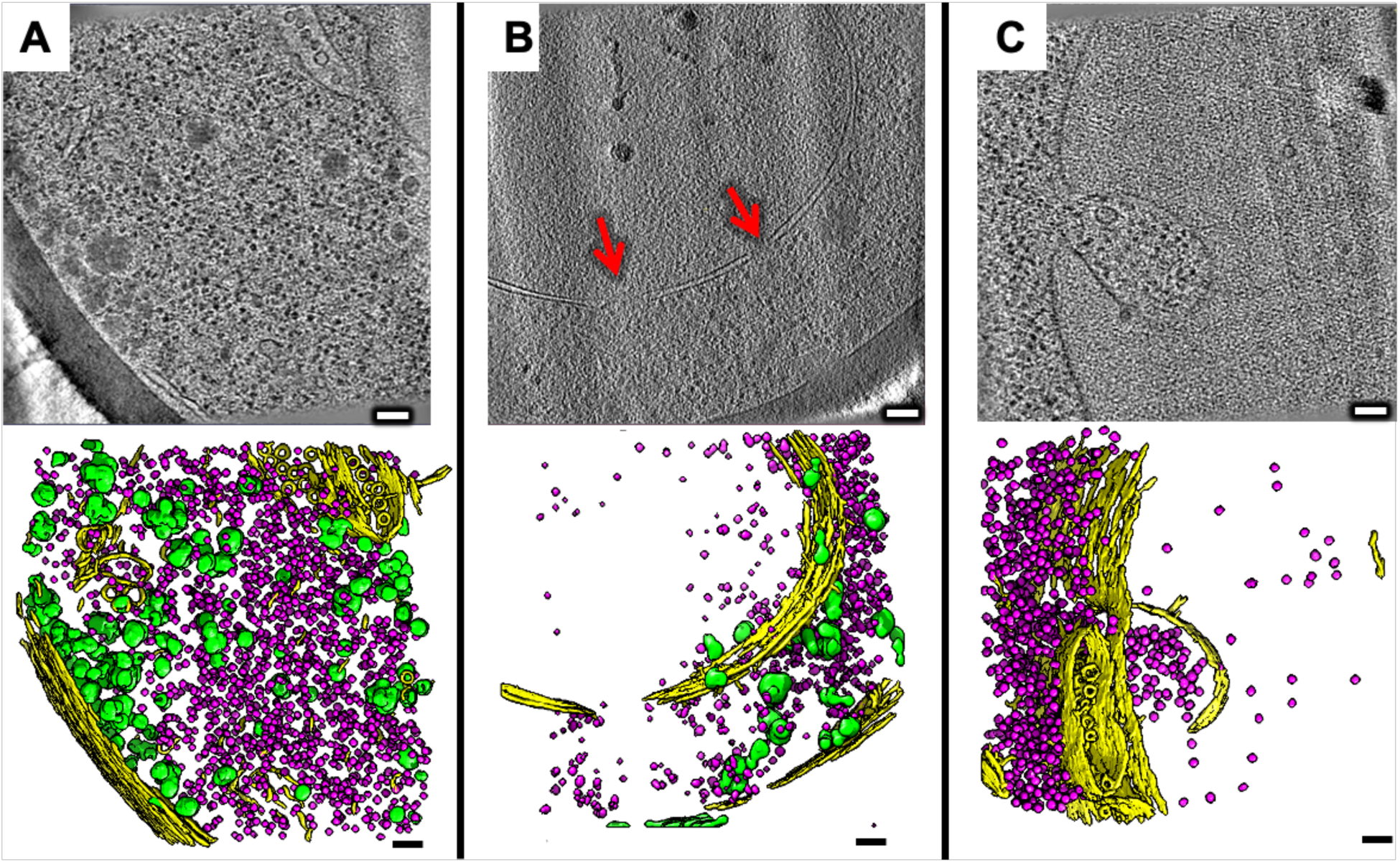
**(A,B)** Slices through representative cryoET tomograms collected from regions with diffuse fluorescence, and **(C)** an area with virtually no fluorescence, and corresponding annotation of features (bottom row) in different colors (green:inclusions containing Hsp104-GFP; yellow:cell wall and membranes, including a multivesicular body in (A) and (C) and nuclear membrane in (B); pink:ribosomes). The red arrows highlight holes in the nuclear membrane, corresponding to nuclear pores. Scale bars: 100 nm.

## DISCUSSION

The goal of visualizing entire cellular atlases for cultured cells and tissues across resolutions is propelling the rapid development of cryoFIB technologies^19^ and correlative approaches with multiple imaging techniques in structural biology. These advances are accelerating our ability to visualize intact cellular and subcellular structures in their native context to better understand their inter-relationships and the structural mechanisms underlying their functions. The formation of the Hsp104-GFP-positive clusters we observed here or similar structures may underlie the reduction in cytotoxic effects from protein misfolding by chaperones^20-22^ and may be central to the reversibility of protein aggregation in heat-shocked yeast when cells are returned to normal temperatures^23^. Indeed, Hsp104 is known to solubilize some protein aggregates^24^ and to modulate prion propagation^25^. More research is needed to determine whether the formation of these granules or their clustering is mediated through phase separation^26^. Furthermore, the workflow we present could also be applied to visualize the structural organization of prion aggregates with heterogenous compositions in yeast^27^ at nanometer resolution and their spatial distributions with respect to other subcellular components to understand possible differences underlying the mechanisms of toxic variants^28^.

While the examples above focused on demonstrating our guided workflow for Hsp104-GFP-positive inclusions at multiple scales (**Supplementary Movie 2**), many additional subcellular components were captured in our 3D “mill and view” cryoFIB-SEM and cryoET data such as vacuoles, lipid droplets, nuclei, ribosomes, mitochondria, and MVBs, among others (**Supplementary Movie 3**), offering ample opportunity for observation-based discoveries. Indeed, we demonstrate that volume “mill and view” can directly visualize multiple vitrified cells at the same time at subcellular resolution, allowing for large-scale, whole-cell imaging and quantification studies of cellular phenotypes under different conditions. Here, we have applied this approach to multiple grids from different cultures of yeast cells, demonstrating the pipeline’s reproducibility. Furthermore, while most institutions may not have all the instruments and expertise for the described experiments, our study demonstrates the feasibility of transferring the fragile milled lamellae across facilities thousands of miles apart (see Methods). These integrated, multi-scale imaging techniques could be applied to a wide variety of cell types to probe cellular and subcellular structures under normal, pathological, and treatment conditions.

## ACKNOWLEDGEMENTS

We thank Profs. Leslie Thompson and William Mobley for many helpful discussions, and Drs. Mei-Kuang Chen and Michael F. Schmid for their feedback on the manuscript, as well as Dr. David Mastronarde for extensive advice (and debugging) in using the latest 3D-CTF correction tools in IMOD.

## AUTHOR CONTRIBUTIONS

This project was conceived by WC with advice from JF on the choice of benchmark samples prepared by EMS and technical guidance for confocal and FIB-SEM microscopy experiments from VG and KC. The CACM experiments were carried out by GHW, PGM and VG. The cryoFIB-SEM experiments were carried out by DM, JM, GHW, PGM, CH, MEB and AKRLJ. The cryoET experiments were carried out by GHW and PGM. Tomographic reconstruction, annotation, and subtomogram averaging were done by JGGM with feedback from GHW. Movie production and editing was done by GHW, CH, VG, KC, and JGGM. The figures were prepared by JGGM and GHW. The manuscript was written mainly by JGGM and WC, with major inputs from GHW, KC, CH & PGM, and feedback from all authors.

## DATA AVAILABILITY AND ACCESSION NUMBERS

Representative tomogram containing an Hsp104-GFP-positive aggregate densities: EMD-21953. Raw CACM, cryoFIB-SEM, and cryoET data necessary to replicate our analyses can be made available upon request.

## STAR METHODS

### Engineering of yeast expressing GFP-tagged Hsp104 and cell culture

Endogenously tagged Hsp104-GFP (S65T) yeast from the Yeast GFP Clone Collection^29^ were grown in YPD medium to an OD_600_ of 0.4. The culture was then incubated at 37 °C for 60 min before vitrification on cryoEM grids for the heat-shocked batch and at room temperature (28 °C) for control cells.

### Sample preparation, vitrification, and pre-screening

Yeast cells were vitrified on Quantifoil 200 mesh gold TEM grids with R2/2 holey carbon film by plunge freezing using a Leica EMGP (5 s back blotting) and clipped into ThermoFisher FIB AutoGrid rings (P/N 1205101) modified for cryoFIB-SEM milling (referred to as “FIB autogrids”). The clipped grids containing Hsp104-GFP yeast were pre-screened using Talos Arctica transmission electron microscope (ThermoFisher) operated at 200 kV to evaluate grid integrity and ice thickness, and to identify suitable cells for subsequent experiments.

### ZEISS cryoAiryscan confocal microscopy screening and subcellular fluorescence target localization

For each EM grid containing vitrified samples, the entire grid was imaged on a Zeiss LSM800 cryoAiryscan microscope equipped with the Linkam CMS196M cryo-stage (maintained at −196 °C) using the 5x EC Plan-Neofluar objective lens (NA 0.16) to acquire low magnification overview images and assess grid quality, sample flatness, and gross ice contamination. Samples were simultaneously imaged in transmitted and fluorescence modes. This allowed us to evaluate the density and distribution of the sample and ensure that the side of the grid containing the cells were cell-side up (facing the objective lens) while providing an overview image for later correlation and re-localization. A coarse z-stack in the reflected-light channel at low magnification (5x) allowed recognition of the “notch”, a directional machined cutout on the FIB autogrid used as a landmark to orient the grid and identify targets of interest in optimal areas for subsequent cryoFIB-SEM experiments^30^. After assessing grid and sample conditions, we restricted imaging to the central region of the grid (most accessible with the gallium ion beam) at intermediate magnification in transmitted and reflected light, as well as Airyscan fluorescence to localize the targets of interest. Once we found suitable targets, we recorded their positions as ROIs in the Zen Blue (Zeiss imaging software). Next, the 100x LD EC Epiplan-Neofluar objective lens (NA 0.75) with a 4.1mm working distance was used to collect 3D confocal z-stacks through the specimen using the Airyscan detector. This Airyscan capability provided high signal-to-noise, high-resolution images using long working-distance air immersion optics under cryo conditions^31^, which facilitated the 3D localization of the targets during correlation across imaging techniques. A coordinate-based overview map with images overlaid as target locations was created using the ZEN Connect module of Zen Blue (Zeiss) and the data was transferred to the cryoFIB-SEM for correlation and target localization. Following Airyscan imaging, the cryo-stage was dismounted from the instrument, and the samples were cryogenically transferred under cryogenic conditions out of the cryo-stage in preparation for cryoFIB-SEM. All experiments up to this point were carried out at Stanford University and Stanford Linear Accelerator Center (SLAC), National Accelerator Laboratory (Stanford-SLAC).

### Cryo focused ion beam and scanning electron microscopy (cryoFIB-SEM)

The clipped FIB AutoGrids with screened frozen samples were then shipped in a Dry Shipper container (CXR100, 3.7 L, Worthington Industries) from Stanford-SLAC to the cryoFIB-SEM imaging facilities at the Massachusetts Institute of Technology (MIT) via FedEx one-day shipping to proceed with subsequent experiments. The specimen grids were then mounted into the FIB-SEM cryo-shuttle (pre-tilted autogrid holder with cryo-shield) under liquid nitrogen in the Leica EM VCM. This shuttle equipped with a sample cryo-shield was then loaded into the VCT500 for cryogenic transfer under vacuum into the chamber of the Zeiss Crossbeam 540 FIB-SEM. The samples were inserted through an airlock on a transfer rod to minimize contamination in the FIB-SEM sample vacuum chamber. Prior to loading of a sample, the stage and anti-contaminator (located within the FIB-SEM sample chamber) were pre-cooled to −150°C and −180°C, respectively, and allowed to equilibrate for 20-30 mins.

Following transfer, the grids were coated with a ∼2-3 μm-thick layer of platinum using the gas injection system (GIS). This helps to reduce charging of and damage to beam-sensitive samples during imaging and milling. For coating, the Pt-GIS was set to 25 °C and the stage was moved to −2 mm from the FIB/SEM coincidence point, before coating for ∼1-2 min. After coating, the grids were screened using the SEM and the FIB. The SEM was used to screen the grids at 538X (0.21 μm/pixel), 2.0 KV, 3.2 pA beam current using the SESI detector to collect secondary electrons and the FIB at ∼273X (409 nm/pixel), 30 kV, 50 pA beam current using the InLens secondary electron detector. The fluorescence maps from the cryoAiryscan were loaded into the ZEN Connect software (Zeiss) and aligned to these SEM and FIB images, allowing for navigation and target selection. The InLens detector allowed for higher-resolution and higher-contrast imaging of unstained samples^32^. Since the samples are very sensitive to gallium ion beam damage, we only used the FIB to image for beam alignment.

### Coarse targeting in z with CACM correlation

The ability to recognize ROIs in all three imaging modalities is helpful for fast and accurate correlation. Features that can be localized in z are particularly valuable, because they can guide where to stop milling. In our case, the pleomorphic clusters correlating with puncta were visible in MAV and low-mag EM images. The z-interval between CACM optical sections was ∼0.42 μm. As we imaged pre-selected ROIs in z-stacks, we used the z-interval and determined the number of z-slices from the top surface of yeast cells before arriving at bright puncta corresponding to Hsp104-GFP signals. Though the cryoFIB-SEM milling slices were much thinner than the z-resolution of CACM, the latter technique provided an informed estimate for how much material to remove during coarse milling. Note that the milling angle was not accounted for during the coarse targeting in z.

The overview images taken using the SEM and FIB in combination with the correlated cryoAiryscan data were used to target ROIs using Zen Connect software. The stage was moved to the center of the ROI and tilted to ∼15-17° to allow for a milling angle of ∼6-8° relative to the grid. Coarse milling was carried out at a pixel size of 39.19 nm, 30 kV, and 700 pA. For monitoring the milling, the SEM was used with a pixel size of 20-50 nm while the voltage and beam currents were 2 kV and 25 pA, respectively.

### Refinement of z-targeting using cryoFIB-SEM “mill and view”

We used “mill and view” (MAV) to further refine the lamella milling boundaries in z, and to obtain contextual 3D information about the regions of the sample that are removed during lamella generation. In general, MAV uses a FIB to remove a thin layer of sample, as thin as ∼3 nm^33^, and images the milled surface with SEM before removing the next thin layer. In our workflow, this procedure can prevent both insufficient and over-milling so long as 1) the puncta are not ablated by coarse milling (this becomes increasingly challenging the smaller the target, given the large step size in CACM stacks), 2) the targets have a depth larger than the MAV slicing interval, and 3) densities corresponding to the targets are recognizable in the cryoFIB-SEM images. MAV records general contextual information of the material removed, allowing us to evaluate cellular features and their positions with respect to ROIs. MAV imaging may find useful applications in large-scale, intermediate-resolution phenotype analyses of vitrified cells and tissues under different disease and treatment conditions.

The MAV SEM imaging settings were 2.96 KX (pixel size=12.56 nm), 2.00 kV, 20 pA, using line integration for noise reduction and an inLens detector, with 30 kV and 50 pA for the focused ion beam and a z step size of 30 nm. The MAV images were aligned using linear stack alignment with SIFT (Fiji). Subsequently, the images were processed using de-striping, CLAHE, and non-local means filters (ORS Dragonfly) to enhance the contrast. Cellular features were manually segmented using Amira^34^ and visualized using UCSF Chimera^35^.

### Final lamella generation

After coarse milling and MAV of the top surface of an ROI, we also milled away material from the bottom side to create the final lamella. We set the FIB to 30 kV and 700 pA for coarse milling to create ∼2 µm-thick lamella, reduced the current to ∼300 pA to thin the lamella to ∼1 µm, and to ∼100 pA to create the final ∼290-500 nm thick lamellae. Finally, we “polished” both sides of the lamellae with a lower current (50 pA).

### Correlating cryoAiryscan and FIB/SEM images with cryoET views

The polished lamellae were obtained with correlative coordinates between the CACM and cryoFIB-SEM images that revealed the positions of aggregates containing Hsp104-GFP. CryoSEM images of polished lamellae show fewer details than MAV images because they are taken with a lower electron dose to minimize electron damage, preserving as much information as possible for subsequent cryoET data collection; however, large aggregates containing Hsp104-GFP are identifiable in low-mag cryoEM screening images. The final lamellae were shipped in a Dry Shipper container (CXR100, 3.7 L, Worthington Industries) from the MIT to the cryoET imaging facilities at Stanford-SLAC via FedEx one-day shipping to proceed with tilt series collection. The fragile samples appeared to be well-preserved through this transportation process, demonstrating that the multi-scale correlative imaging can be carried out with instrumentation in multiple facilities.

### CryoET tilt series data collection, tomographic reconstruction, annotation, and subtomogram averaging

We collected tilt series of lamellae containing vitrified yeast cells with (n=17) and without (n=34) heat shock using a Titan Krios electron microscope (ThermoFisher) operated at 300 kV in low-dose mode using SerialEM software^36^. At each tilt angle, we recorded “movies’’ with 5 frames each using a K2 camera (Gatan). The tilt series were collected bi-directionally from 25° to −60°, and then from 26° to +60° with a tilt step of 1°, target defocus of −8 μm and a cumulative dose of ∼100 e/Å^2^. The images were motion-corrected with MotionCor2 software^37^. We used the IMOD software package^38^ for standard weighted-back projection tomographic reconstruction after patch-tracking alignment. Within each tilt series, unusable images (large drift, excessive ice contamination, etc.) were removed prior to tilt series alignment. The data were contrast transfer function (CTF) corrected with IMOD’s newly implemented 3D-CTF correction algorithm. Parallel reconstructions using a SIRT-like filter were computed for visualization purposes using heavily binned data (shrink factor=4) and post-reconstruction filters (lowpass, highpass, normalization, and thresholding at 3 standard deviations away from the mean). We carried out tomographic annotation of different features in the binned-by-4 tomograms using EMAN2’s semi-automated 2D neural network-based pipeline^37^, and performed manual clean-up of false positives in UCSF Chimera, which we also used to display all annotated densities and prepare the cryoET movies.

## SUPPLEMENTARY DATA

**Supplementary Figure 1.**
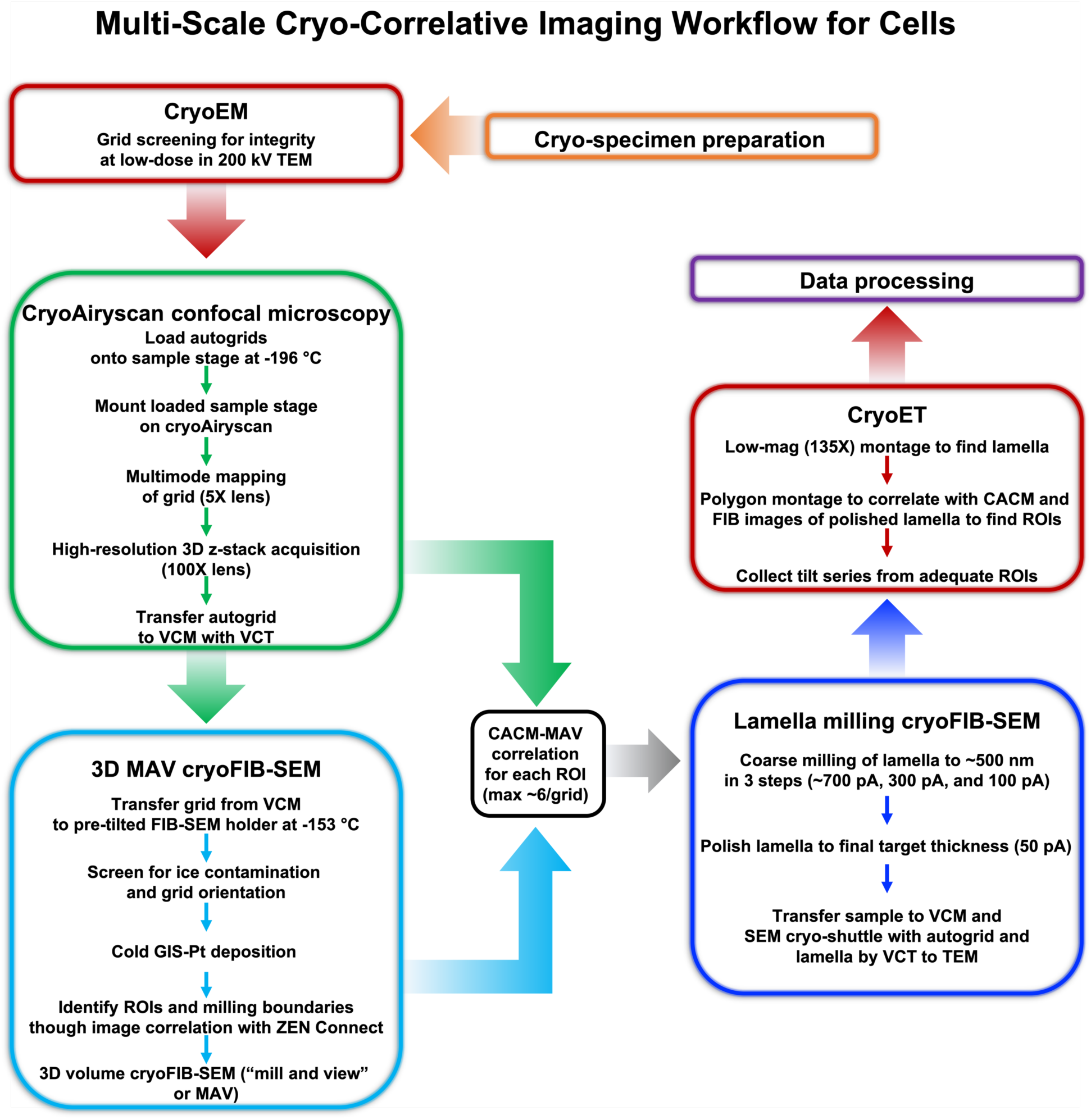
Overview schematic of multi-scale cryo-correlative imaging workflow for cells; kV:kilo Volt; TEM:transmission electron microscope; VCM:; VCT:; GIS: gas injection system; Pt:platinum; ROI:region of interest; CACM:cryoAiryscan confocal microscopy; MAV:mill and view.

**Supplementary Movie 1**

Cartoon depiction of vitrified yeast cells containing a fluorescently labelled inclusion and imaged by “mill and view” (MAV) cryoFIB-SEM, followed by lamella production, and cryoET imaging.

**Supplementary Movie 2**

This movie illustrates our entire multi-scale 3D cryo-correlative workflow with an experimental example of data from vitrified heat-shocked yeast cells expressing the endogenous protein chaperone Hsp104 bearing a C-terminal GFP fusion tag, imaged with CACM, followed by cryoFIB-SEM MAV and lamella generation, and ending with cryoET and annotation of features.

**Supplementary Movie 3**

Superposition of 3D data and corresponding annotations derived from the visualization of frozen-hydrated heat-shocked yeast cells expressing Hsp104-GFP with three different imaging instruments and modalities. The data include a representative CACM volume stack (fluorescent green signals), cryoFIB-SEM MAV volume stack, and cryoET tomogram.

